# Massive outbreak of Influenza A H5N1 in elephant seals at Península Valdés, Argentina: increased evidence for mammal-to-mammal transmission

**DOI:** 10.1101/2024.05.31.596774

**Authors:** Marcela Uhart, Ralph E. T. Vanstreels, Martha I. Nelson, Valeria Olivera, Julieta Campagna, Victoria Zavattieri, Philippe Lemey, Claudio Campagna, Valeria Falabella, Agustina Rimondi

## Abstract

H5N1 high pathogenicity avian influenza (HPAI) viruses of the clade 2.3.4.4b have killed thousands of marine mammals in South America since 2022. In October 2023, following outbreaks in sea lions in Argentina, we recorded unprecedented mass mortality (∼17,000 individuals) in southern elephant seals (*Mirounga leonina*) at Península Valdés. Seal pups were disproportionately affected. Adult seals departed early, disrupting social and breeding structure. Frequent interactions with sea lions and scavenging by seagulls were observed. Deaths of terns concurred with seals but peaked weeks later. HPAI H5N1 was confirmed in seals and terns. Moreover, genomic characterization showed viruses from pinnipeds and terns in Argentina form a distinct clade with marine mammal viruses from Peru, Chile and Brazil. These mammal-clade viruses share an identical set of mammalian adaptation mutations which are notably also found in the terns. Our combined ecological and phylogenetic data support mammal-to-mammal transmission and occasional mammal-to-bird spillover. To our knowledge, this is the first multinational transmission of H5N1 viruses in mammals ever observed globally. The implication that H5N1 viruses are becoming more evolutionary flexible and adapting to mammals in new ways could have global consequences for wildlife, humans, and/or livestock.

## INTRODUCTION

The emergence of H5N1 high pathogenicity avian influenza (HPAI) viruses from clade 2.3.4.4b in 2020 triggered numerous outbreaks in wildlife worldwide^1^. In Europe and southern Africa, impacts to wildlife were particularly severe in seabird colonies, with losses in the tens of thousands^2–5^. In 2021–2022, these H5N1 HPAI 2.3.4.4b viruses spread to North America, further impacting wildlife, especially waterbirds and birds of prey^6^ and reassorting with endemic strains^7,8^. The virus then spread to South America in 2022 via multiple introductions^9,10^, causing unprecedented large-scale mortality of seabirds, with an estimated death toll surpassing 650,000 individuals^10–14^.

H5N1 HPAI sporadically caused mortality of pinnipeds and cetaceans in Europe^15,16^ and North America^17–19^, but it was only upon reaching the Pacific coast of South America that the virus demonstrated an ability to cause large-scale mortality in marine mammals^10,20^. More than 30,000 South American sea lions (*Otaria byronia*) died as H5N1 virus spread along the coast of Peru and Chile in 2022–2023, with porpoises, dolphins and otters also being affected in smaller numbers^10,12–14,20–22^. Following the southward spread along the Pacific coast of South America, H5N1 HPAI viruses were detected in sea lions at the southern tip of Chile in June 2023^22^. By early August, the virus was detected for the first time on the Atlantic coast, in a sea lion rookery off southernmost Argentina. Then, over the following weeks, the virus spread rapidly northward along Argentina’s Atlantic coast, killing hundreds of sea lions along Argentina’s shores^23^, eventually reaching Uruguay^24^ and southern Brazil^25^.

Shortly thereafter, in October 2023, we recorded unprecedented mass mortality in southern elephant seals (*Mirounga leonina*) at Península Valdés in central Patagonia, Argentina, with an estimated death toll surpassing 17,000 individuals^26^. In this study, we present epidemiological data and full genome characterization of H5N1 clade 2.3.4.4b viruses associated with the outbreak in elephant seals and with concurrent tern mortality. We analyze data from the Península Valdés event and prior reports to investigate potential pathways of H5N1 virus transmission among marine mammals and birds in South America, and document a rapidly spreading H5N1 marine mammal clade carrying mammalian adaptation mutations of potential public health concern.

## RESULTS

### Elephant seal mortality at Punta Delgada breeding colony, Península Valdés

On 10-Oct-2023, we surveyed the breeding colony at Punta Delgada (Península Valdés, Argentina), and counted 218 living and 570 dead pups (including weaners) (Table 1, Figure 1A). This represented more than 70-fold increase in pup mortality rate compared to the prior year (71% in 2023 vs. 1% in 2022). By 13-Nov-2023, only 38 pups survived (95% mortality). At least 35 subadult/adult seal carcasses were recorded in the area, whereas in previous years even a single dead adult seal was a rare sighting. Retrospective inquiries suggest that navy personnel at the Punta Delgada lighthouse first observed signs of unusual mortality around 25-Sep-2023, but the finding was not reported at that time. No unusual mortality was seen in juveniles, which began gathering in usual numbers in November (Table 1).

**Figure 1.**
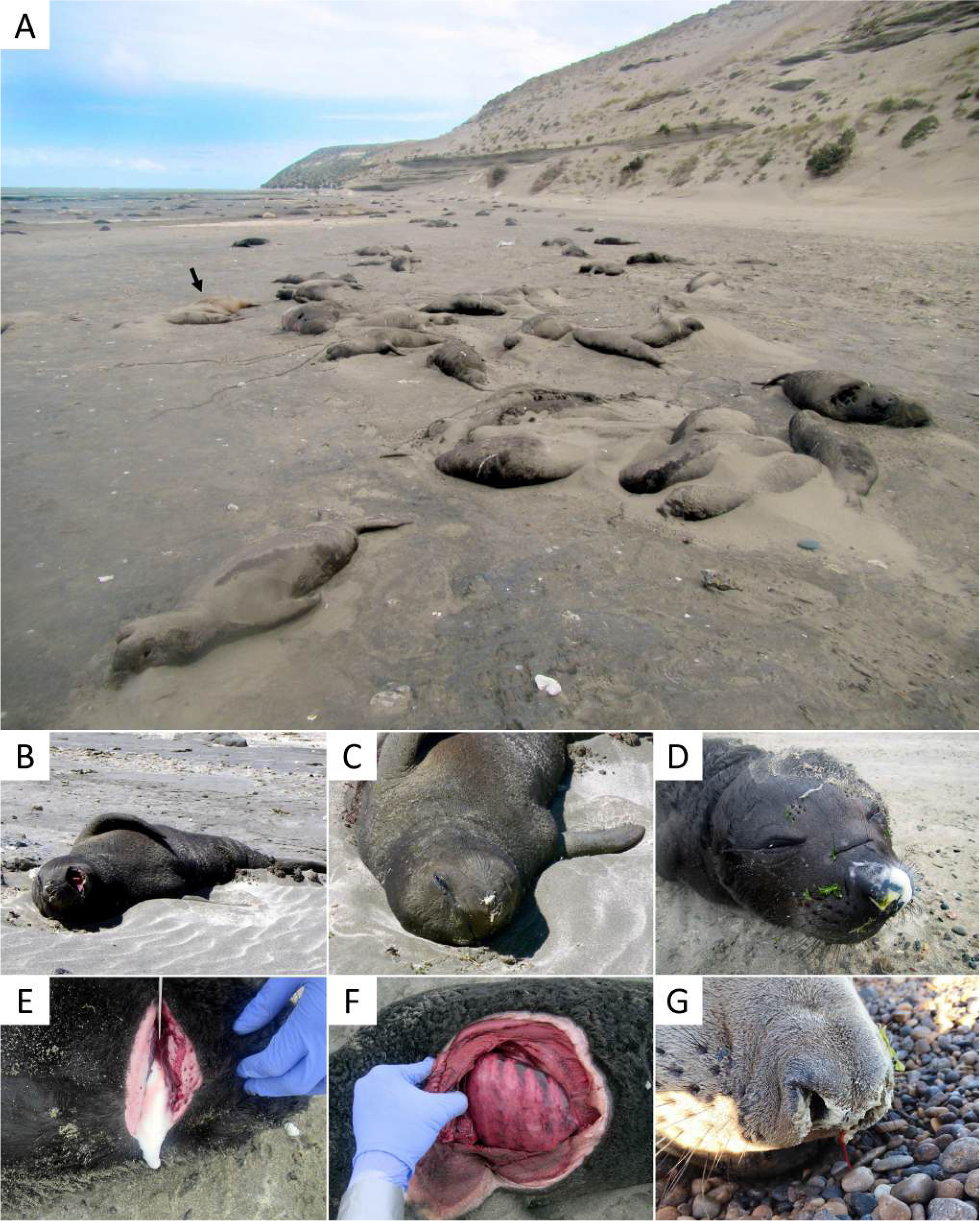
Mass mortality, clinical signs and post-mortem findings of elephant seals at Punta Delgada (Península Valdés, Argentina) during an outbreak of H5N1 HPAI. A) Hundreds of elephant seal pup carcasses accumulated along the high tide line of the beach at Punta Delgada; a sea lion carcass (arrow) and patchily distributed living elephant seals (far background behind the arrow) are also visible. B) Pup presenting with labored breathing and foamy nasal discharge. C) Pup presenting with open mouth breathing and tremors/twitching. D, E) Abundant white foam on the snout and draining from the trachea of a dead pup. G) Markedly heterogeneous and congested lung surface in a dead pup. H) Bloody and mucous nasal discharge in a dead subadult male.

**Table 1.**
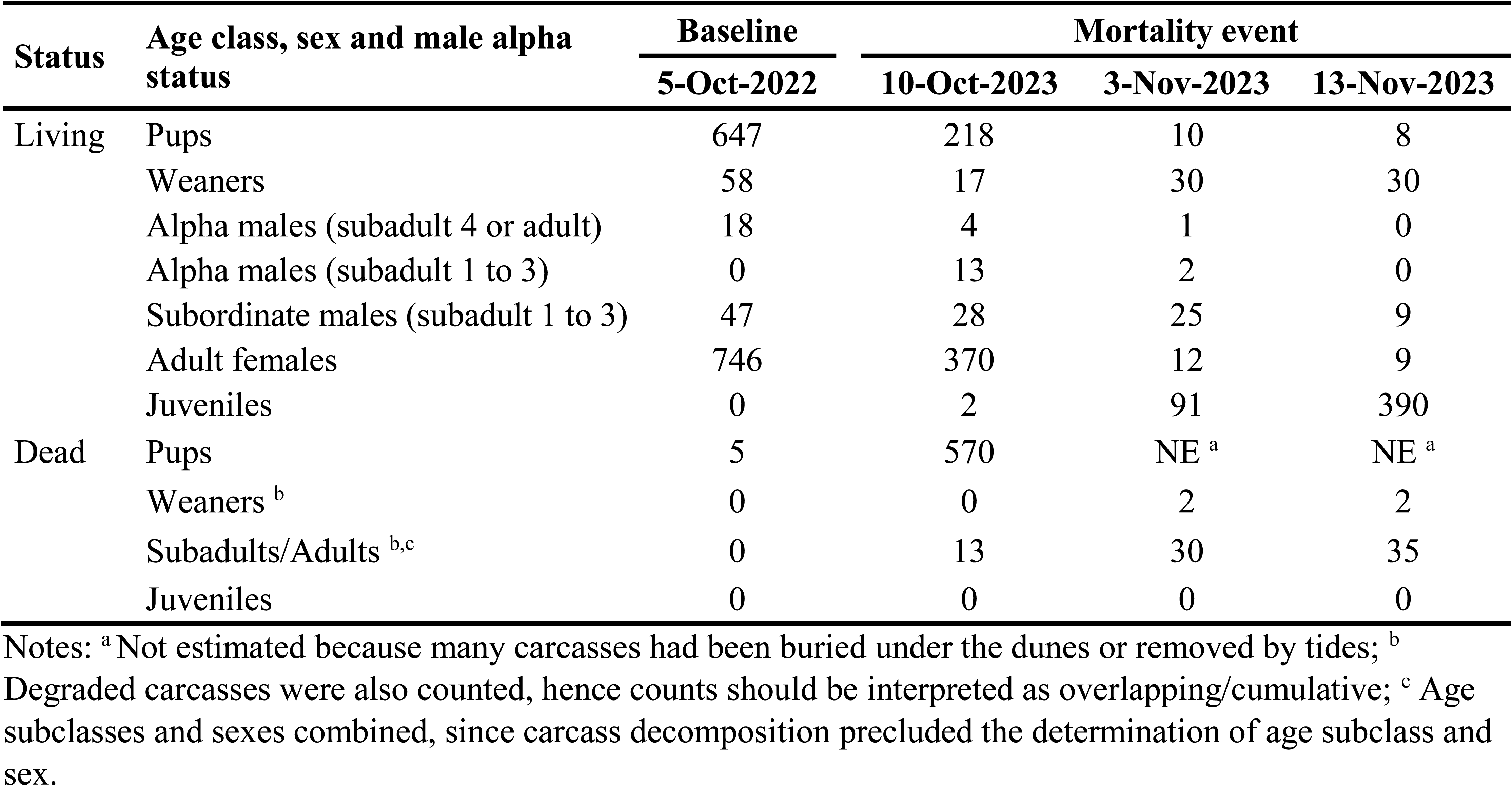
Number of living and dead southern elephant seals (*Mirounga leonina*) at Punta Delgada breeding colony (Península Valdés, Argentina) during the 2022 season (baseline) and the 2023 mortality event.

| Status | Age class, sex and male alpha status | Baseline | Mortality event |  |  |
| --- | --- | --- | --- | --- | --- |
|  |  | 5-Oct-2022 | 10-Oct-2023 | 3-Nov-2023 | 13-Nov-2023 |
| Living | Pups | 647 | 218 | 10 | 8 |
|  | Weaners | 58 | 17 | 30 | 30 |
|  | Alpha males (subadult 4 or adult) | 18 | 4 | 1 | 0 |
|  | Alpha males (subadult 1 to 3) | 0 | 13 | 2 | 0 |
|  | Subordinate males (subadult 1 to 3) | 47 | 28 | 25 | 9 |
|  | Adult females | 746 | 370 | 12 | 9 |
|  | Juveniles | 0 | 2 | 91 | 390 |
| Dead | Pups | 5 | 570 | NE <sup>a</sup> | NE <sup>a</sup> |
|  | Weaners <sup>b</sup> | 0 | 0 | 2 | 2 |
|  | Subadults/Adults <sup>b,c</sup> | 0 | 13 | 30 | 35 |
|  | Juveniles | 0 | 0 | 0 | 0 |

Notes: <sup>a</sup> Not estimated because many carcasses had been buried under the dunes or removed by tides; <sup>b</sup> Degraded carcasses were also counted, hence counts should be interpreted as overlapping/cumulative; <sup>c</sup> Age subclasses and sexes combined, since carcass decomposition precluded the determination of age subclass and sex.

The mortality event led to drastic changes in the elephant seal social structure (Table 1), with a progressive replacement of mature alpha males by subadults and rapid decline in numbers of breeding females. This manifested as a patchy distribution of seals with scattered females without pups as well as abandoned and sick pups. By 13-Nov-2023, all breeding structure was dissolved. There were no harems, only 9 males (all subadults not associated with females) and 9 females (8 with pup and 1 pupless) amidst carcasses of elephant seals (Supplementary Figures 1A and 1B).

The absence of large alpha male elephant seals to chase away perceived intruders resulted in a larger number of South American sea lions commingling or dead among breeding elephant seals at Punta Delgada (Table 2). This prompted agonistic interactions with nursing elephant seal mothers (Supplementary Figure 1C) and attempts at sexual interactions with pups (Supplementary Figure 1D). Other interspecies interactions included the scavenging of elephant seal carcasses by kelp gulls (*Larus dominicanus*) (Supplementary Figure 1E) and the presence of living and dead South American terns (*Sterna hirundinacea*) amidst elephant seal carcasses (Supplementary Figure 1F). Some terns showed neurological signs of disorientation, decreased fear response and difficulty/inability to fly, and were not in social groups as would be expected. The tern death toll increased over time to almost 400 dead birds (Table 2).

**Table 2.**
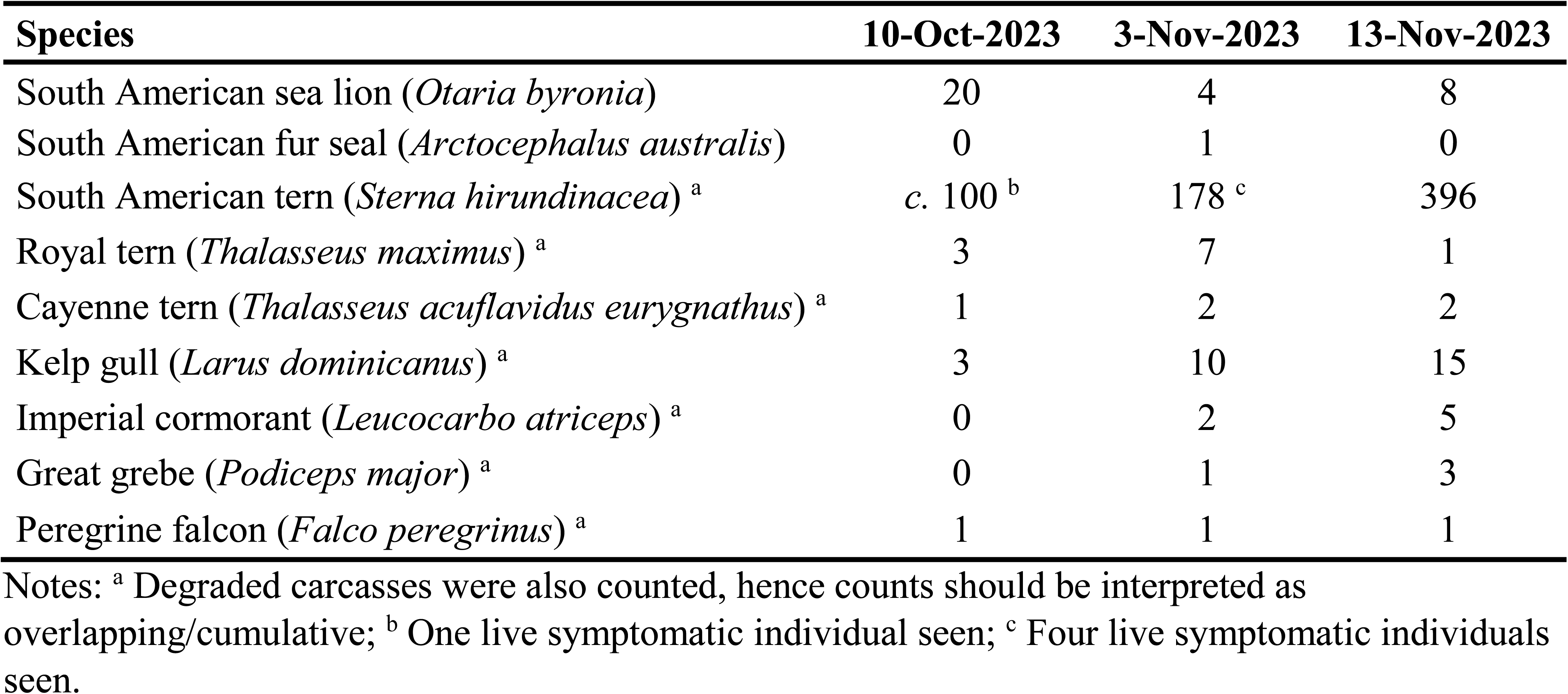
Estimated number of dead individuals of other pinniped species and seabirds at Punta Delgada (Península Valdés, Argentina) in 2023, during the elephant seal mortality event.

As per the temporal distribution of events, mortality of elephant seal pups peaked between 25-Sep-2023 and 10-Oct-2023, whereas the majority of terns died about three weeks later, between 3-Nov-2023 and 13-Nov-2023. This temporal delay also occurred in Argentina as a whole, with large-scale mortalities of sea lions (mid-August to late September 2023) and elephant seals (late September to mid-October 2023) preceding the large-scale mortality of terns (early to mid-November 2023).

### Clinical signs and post-mortem findings in elephant seals

Elephant seal pups showing clinical signs consistent with HPAI were seen during all field surveys in October and November 2023. Symptomatic pups were lethargic, had difficulties to roll or galumph, and labored breathing, nasal discharge, repetitive head or flipper movements and tremors (Figures 1B–D, Supplementary File 1). Most symptomatic pups were motherless and alone or close to other abandoned or dead pups. During one field survey, several pups were seen at risk of drowning with the incoming tide (Supplementary File 1). Ill and dead pups ranged in age from newborn to about 3 weeks-old (i.e. about to wean). Some carcasses of freshly deceased pups showed foam or mucous nasal discharge (Figure 1D), and abundant white foam drained from the sectioned trachea of one individual (Figure 1E). It is unclear whether this was due to infection or agonal drowning. The lungs of four pups showed a heterogeneous and congested surface (Figure 1F), draining blood profusely when cut. We did not perform full necropsies due to biosecurity concerns; hence, we did not examine other organs. Following deaths in the breeding areas, several elephant seals hauled out at a second, aberrant location (Golfo Nuevo) in October-December 2023 (Supplementary Figure 2, Supplementary Table 1). Of these, one subadult male died within two days after showing clinical signs consistent with HPAI (tremors, labored breathing, yellowish and blood-stained nasal discharge, hyperthermia; Figure 1G, Supplementary File 1).

### H5N1 HPAI viruses belong to clade 2.3.4.4b and genotype B3.2

We tested swab samples from four elephant seal pups, five South American terns and two royal terns (pooled according to sample type) and an additional pool containing all samples from a sixth South American tern from Punta Delgada. Another pool containing all samples from a dead subadult male elephant seal from Golfo Nuevo was also analyzed. All pools were positive for the matrix gene of influenza A virus. Pools from elephant seals were further tested for H5 clade 2.3.4.4b and were also positive (Supplementary Table 2). We performed whole genome sequencing for: a pool of samples from one South American tern (CH-PD037) from Punta Delgada, brain samples from one elephant seal pup (CH-PD035), one South American tern (CH-PD030) and one royal tern (CH-PD036) from Punta Delgada, and a rectal sample from the symptomatic subadult male elephant seal (CH-PM053) from Golfo Nuevo. We obtained five H5N1 HPAI virus genomes in this study (Supplementary Table 3) and nucleotide sequences were deposited in GenBank (accession numbers PP488310–PP488349).

### Distinct HPAI H5N1 viruses in avian and mammalian hosts

We first compared our H5N1 HPAI viruses in Península Valdés with other strains from South America, North America, and Eurasia during 2021–2023 to confirm that the H5N1 HPAI outbreaks that occurred in Argentina, Brazil, Chile, Peru, Uruguay, and Antarctica from November 2022 to November 2023 all stem from a single introduction of clade 2.3.4.4b genotype B3.2^8^ from North American wild birds into South America (Supplementary Figure 3). B3.2 viruses have a reassortant genotype with four segments from the Eurasian H5 lineage (PA, HA, NA, and MP) and four segments from low pathogenicity avian influenza viruses from the North American lineage (PB2, PB1, NP, and NS). Our analysis shows that all H5N1 HPAI viruses in Argentina have the 4:4 reassortant genotype B3.2, including the five viruses sequenced for this study, six viruses sequenced from our previous report^23^, and 46 viruses from poultry and one wild bird (Andean goose) available in GISAID. However, Argentina’s H5N1 HPAI viruses are not monophyletic (i.e., clustering together as a single Argentina clade, separate from viruses from other countries). Instead, viruses collected from poultry in inland Argentina cluster separately from viruses collected from Argentina’s coastal outbreaks in marine mammals and terns (Figure 2, Supplementary Figures 4–5). Argentina’s poultry viruses are positioned in a clade on the tree that includes (a) Argentina’s earliest detected H5N1 HPAI virus (A/goose/Argentina/389-1/2023; this virus was detected on 11-Feb-2023 and is the only currently available sequence for H5N1 HPAI viruses from inland wild birds in Argentina), (b) poultry viruses from Uruguay and Chile, and (c) some wild bird viruses from Uruguay, Brazil, Chile, and Antarctica. Within this clade, Argentina’s poultry viruses are intermixed with viruses from other locations and wild bird hosts, suggesting frequent virus movement across national borders and spillover between wild birds and poultry.

**Figure 2.**
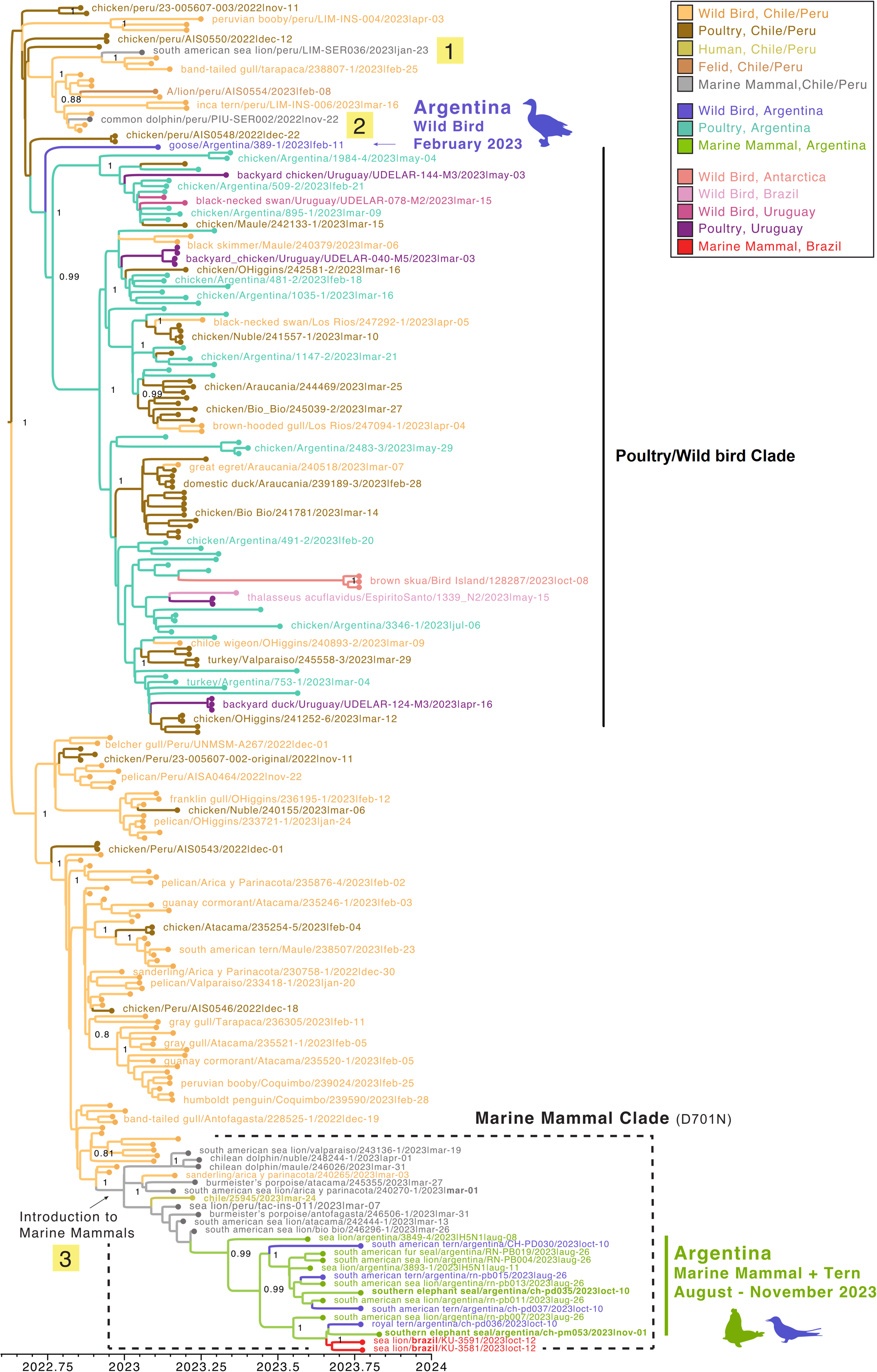
Phylogenetic tree of H5N1 HPAI (2.3.3.4b) viruses in South America. Time-scale MCC tree inferred for the concatenated genome sequences (∼13kb) of 236 H5N1 influenza A viruses (clade 2.3.3.4b) collected in five South American countries (Argentina, Brazil, Chile, Peru, Uruguay) and Antarctica. Three spillover events into marine mammals, including the marine mammal clade, are labeled. Branches are shaded by inferred host species and location (13 categories). Posterior probabilities provided for key nodes. Tip labels provided for all mammalian viruses and a selection of avian viruses.

Conversely, the vast majority of wild bird viruses from Peru and Chile are positioned in a different clade (lower section of the tree in Figure 2). This wild bird clade is closely related (and basal) to a clade of marine mammal viruses collected from four countries (Peru, Chile, Argentina, and Brazil). A quantitative estimate of virus gene flow in the ancestry of our sample (through “Markov jump” counts, Figure 3A) indicates that H5N1 HPAI viruses transmitted ∼3x from wild birds to marine mammals on the Pacific (western) coast of South America. Two wild bird-to-marine mammal transmissions in Peru appear to be dead-end spillover events with no secondary cases (A/common dolphin/Peru/PIU-SER002/2022 and A/South American sea lion/Peru/LIM-SER036/2023). In contrast, a third wild bird-to-marine mammal transmission is associated with a multinational clade of 26 viruses, including 20 from marine mammals in Peru (n = 1), Chile (n = 8), Argentina (n = 9), and Brazil (n = 2). The time-scaled MCC tree estimates that the third wild bird-to-marine mammal transmission event occurred between 24-Nov-2022 and 7-Jan-2023, based on the estimated time to the most recent common ancestor (Figure 2). The multinational marine mammal clade also includes a human case from Chile (A/Chile/25945/2023), a wild bird virus from Chile (A/sanderling/Arica y Parinacota/240265/2023), and four viruses obtained from terns in Argentina (one South American tern from Punta Bermeja in August 2023, one royal tern and two South American terns from Punta Delgada in October 2023) that are closely related to the marine mammal viruses from Argentina. Our results showed that the human, sanderling, and four tern viruses positioned in the marine mammal clade appear to be independent spillover events from marine mammals (Figures 2 and 3A). This is further supported by the fact that these viruses share mutations in PB2 that are associated with mammalian adaptation and are present in viruses forming to the marine mammal clade (Figure 4). Figure 5 summarizes our hypothesized pathway of spread of H5N1 HPAI viruses in South America based on the molecular evidence and the chronology of reported detections.

**Figure 3.**
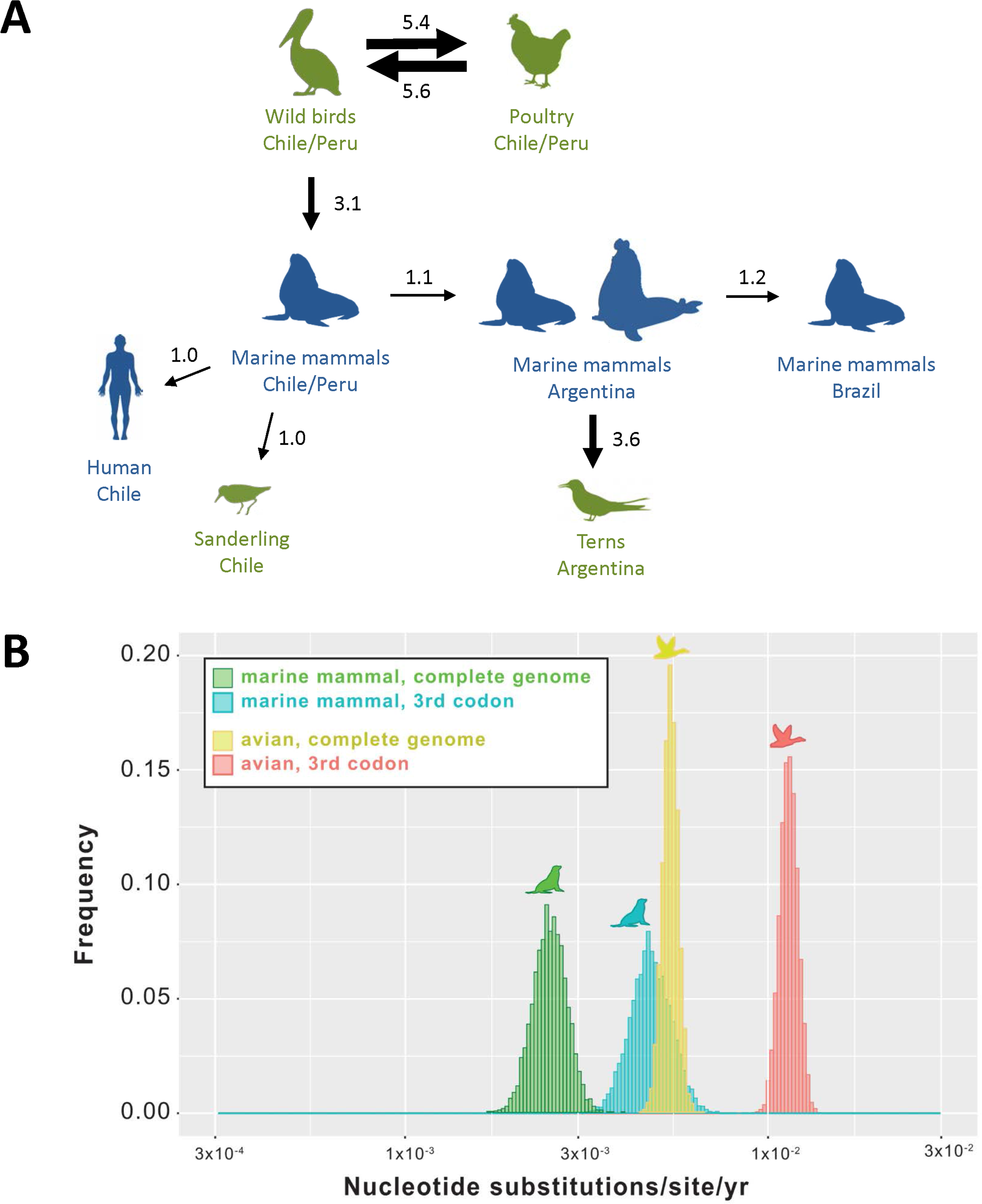
Phylodynamics of H5N1 HPAI (2.3.4.4b) viruses in South American marine mammals and birds. (A) Direction of virus gene flow between locations and hosts, inferred from “Markov jump” counts across the posterior distribution of trees inferred using a Bayesian approach (values under 0.5 excluded). Different host groups are indicated with different colors: avian (green) and mammal (blue). (B) Posterior distributions of evolutionary rates (substitutions per site per year) inferred for the complete virus genome (all positions) and for only the third nucleotide position for H5N1 (2.3.4.4b) in South America, partitioned into two host categories: marine mammal and wild bird/poultry.

**Figure 4.**
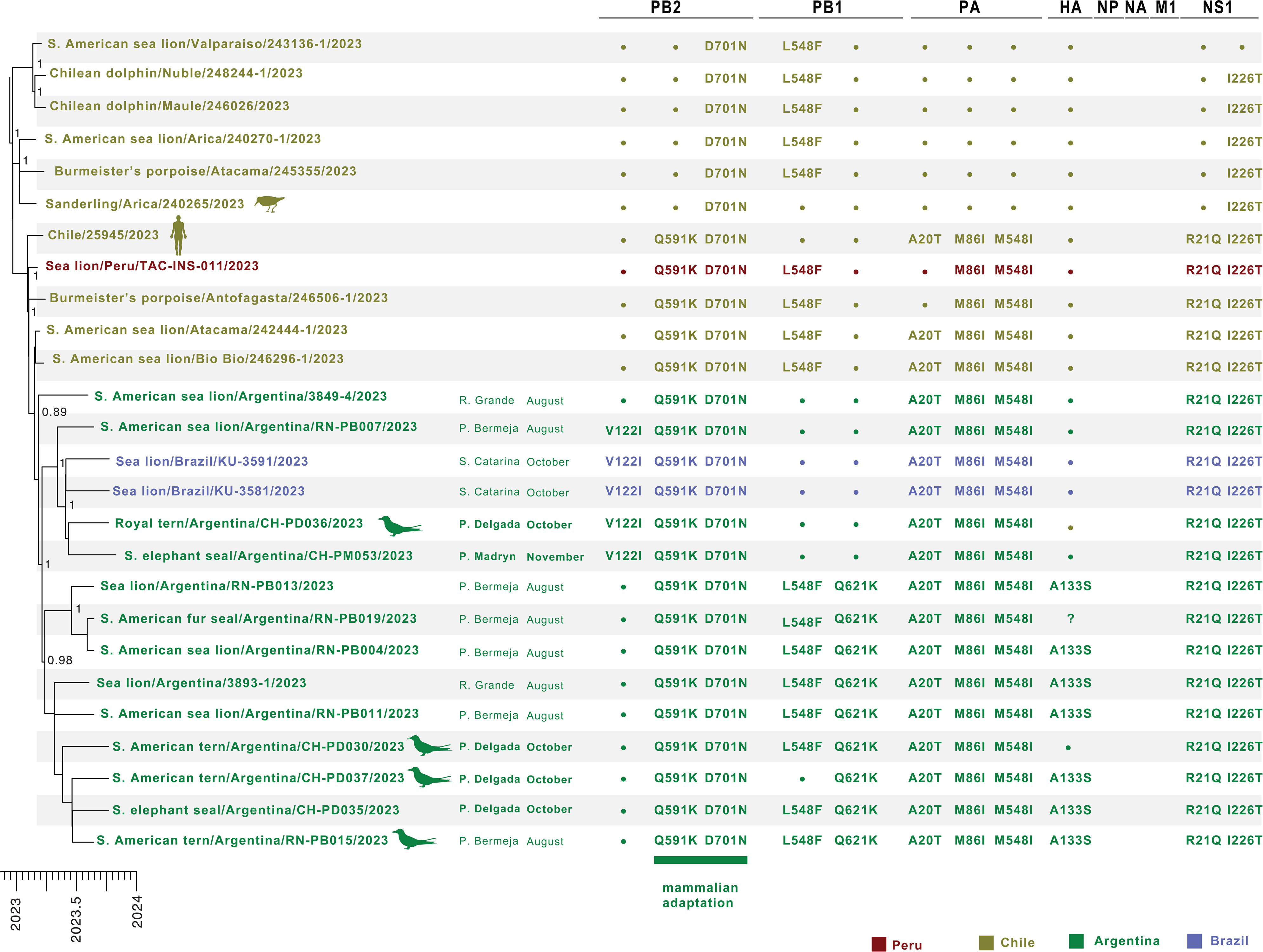
Mutations defining the marine mammal clade of H5N1 HPAI (2.3.4.4b) viruses. Amino acid changes are listed for new mutations that arose in the marine mammal clade of the H5N1 HPAI (2.3.4.4b) viruses that are not observed in any other avian viruses included in this study from South America, mapped against the subsection of the MCC tree with the marine mammal clade (see Figure 2). Virus names and associated mutations are colored by country. Location/month of collection (in 2023) are listed for Argentina and Brazil. A question mark indicates that no sequence data is available at that position for that virus. H5 numbering is used for HA.

**Figure 5.**
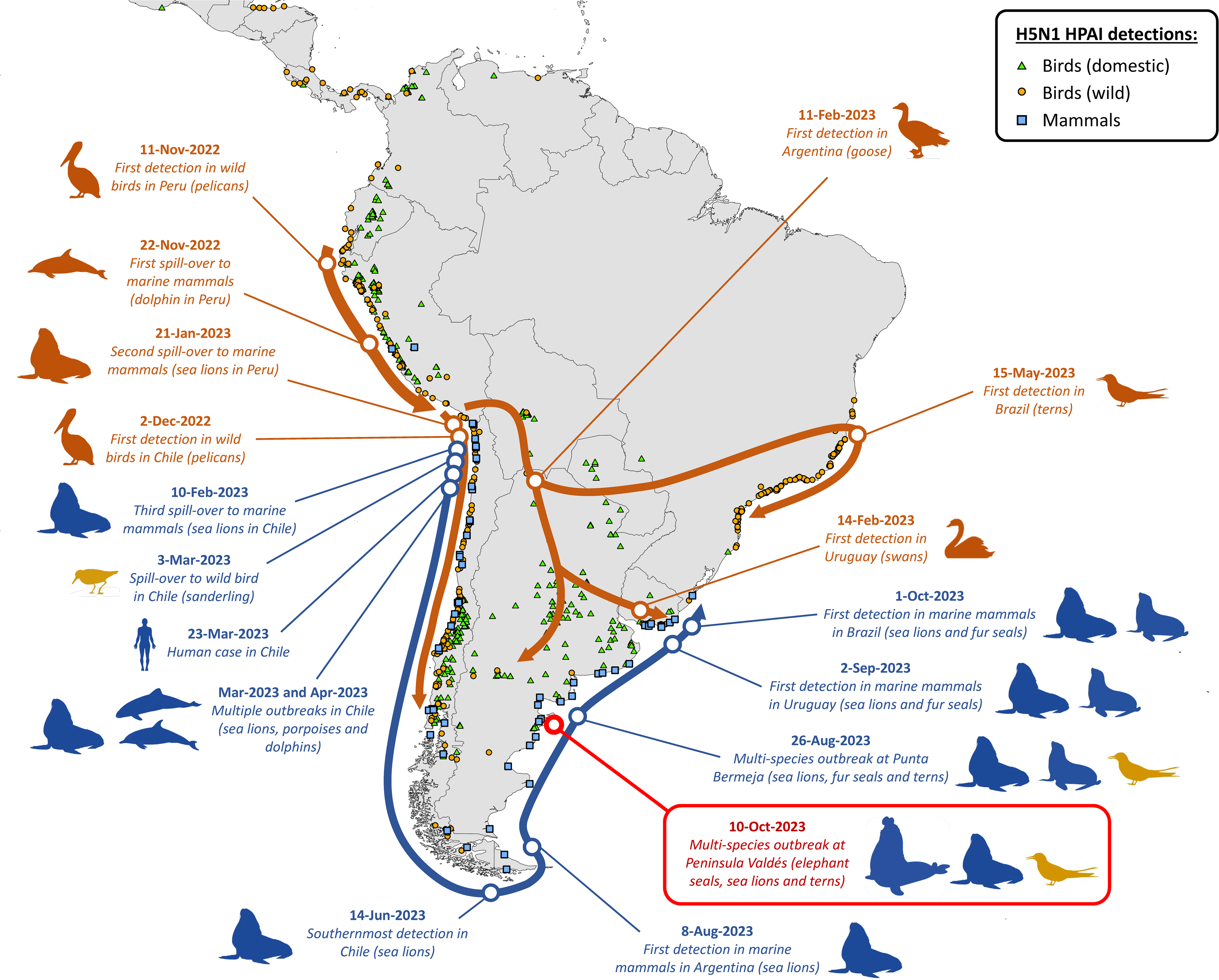
Chronology and hypothesized pathways of spread of H5N1 HPAI (2.3.4.4b) viruses in South America, 2022–2023. H5Nx HPAI detections (1-Sep-2022 to 1-Nov-2023) reported to the World Animal Health Information System (WAHIS/WOAH) are represented with orange circles (wild birds), green triangles (domestic birds) and blue squares (mammals). The location of the outbreak investigated in this study (Península Valdés) is highlighted in red. Arrows represent the timeline of hypothesized pathways of virus spread, as derived from the chronology of detections and our phylodynamic analysis. The pathways of virus spread and significant events of the avian and marine mammal clade viruses are represented in dark orange and dark blue, respectively. Dark yellow represents incidental avian hosts of marine mammal clade viruses (i.e. spillover). Note that virus spread pathways in this figure are intended as a conceptual model and are not geographically precise.

### Lower evolutionary rate of HPAI H5N1 viruses in marine mammals

To account for the possibility that convergent evolution following host-switches could cause marine mammal viruses to artificially cluster on the tree when they do not actually share common ancestry, phylogenies were inferred for: (a) the entire virus genome (∼13kb) (Figure 2) and (b) the third codon position only (Supplementary Figure 6). The similarity of the two trees suggests that the marine mammal clade is not merely an artifact of strong convergent evolution for adaptive mutations in marine mammals following a host-switch, but rather that the marine mammal clade is real (Supplementary Figures 4 and 12). If H5N1 HPAI viruses are transmitting independently in marine mammals across multiple South American countries, a host-specific local clock (HSLC) should be used to accommodate a different rate of evolution. The estimated rate of evolution in the marine mammal clade (human and avian viruses excluded) using a HSLC was ∼2-fold lower (2.5 x 10^-3^; 2.0–3.0 x 10^-3^ 95% HPD) than the avian rate (5.4 x 10^-3^; 4.9–5.9 x10^-3^ 95% HPD), which includes wild birds and poultry but excludes spillovers into mammals (Figure 3B). The marine mammal rate was still ∼2-fold lower compared to birds when only the third codon position was considered (Figure 3B). The eight genome segments showed strong purifying selection in both avian and marine mammal hosts in South America, with dN/dS ratios under 0.3 (Supplementary Figure 7), comparable to previous estimates^27^.

### Global SNP analysis reveals mammal adaptation mutations and suggests two HPAI H5N1 subpopulations during mammal-to-mammal transmission in Argentina

Across the genome, we identified more than 64 amino acid changes in the H5N1 HPAI viruses from Península Valdés when compared viruses from birds and mammals from Argentina, other South American countries, Antarctica, North America (genotype B3.2 from 2022–2023) and the original Goose/Guangdong (Gs/Gd) (Supplementary Table 4). Of the 64 mutations, 18 are potentially associated with increased virulence, transmission or adaptation to mammalian hosts, and fifteen are present in H5N1 viruses from Argentina’s coastal outbreaks in marine mammals and terns but absent in H5N1 (B3.2 genotype) strains from North America and from goose/poultry strains from Argentina (Supplementary Table 4). Of note, eleven of the fifteen common mutations were also present in the human case in Chile (Supplementary Table 5).

Argentina’s marine mammal viruses inherited eight amino acid changes that emerged previously in marine mammals in Chile and Peru that were never seen in H5N1 HPAI viruses circulating in birds in South America and appear to be specific to the marine mammal clade (Figure 4): Q591K and D701N in PB2; L548F in PB1; A20T, M86I, and M548I in PA; and R21Q and I226T in NS1. Almost all mutations (except L548F in PB1) were also present in the two Brazil marine mammal viruses. The conservation of seven amino acid changes across all marine mammal viruses collected from three countries over eight months (Chile, Argentina, Brazil; March through October) further supports the existence of an independent chain of virus transmission among marine mammals, separate from avian transmission chains. In addition to nonsynonymous mutations, four silent mutations in PB1 (A1167T), PA (C1359T), and NP (C669T and T1239C) were found in marine mammal viruses in Argentina that were inherited from marine mammal viruses circulating in Peru and/or Chile (Supplementary Figure 8).

Synonymous and non-synonymous mutations also occurred during H5N1 2.3.4.4b circulation in Argentina, leading to the evolution of two distinct subpopulations defined by specific mutations. The first Argentina subpopulation is defined by a new V122I substitution in PB2 and the loss of the L548F substitution in PB1 (owing to a secondary substitution), and also detected in marine mammals in Brazil (Figure 4). The second Argentina subpopulation is defined by a new Q621K substitution in PB1, which in almost all cases is accompanied by mutation A133S in HA. This A133S substitution in HA that was seen in Argentina in South American terns (n = 2), South American sea lions (n = 4), and an elephant seal (n = 1) (note: this HA region could not be sequenced from the South American fur seal) was not observed in previous H5N1 pinniped outbreaks in South America, North America or Europe, nor in bird outbreaks from South America (Figure 4, Supplementary Table 5). Of note, both subpopulations were found in the terns and elephant seals sampled for this study and in mammalian and avian hosts in a multi-species outbreak at Punta Bermeja (∼260 km north of Punta Delgada) in August 2023^23^, but not in the first H5N1 HPAI detection in Argentina, nor in any of the viruses from poultry in this country (Figure 4, Supplementary Table 5).

## DISCUSSION

Since 2020, the world has witnessed an unprecedented global epizootic of H5N1 clade 2.3.4.4b viruses with a catastrophic ecological impact on wildlife species, including pinnipeds. Although H5N1 HPAI viruses were previously implicated in mortalities of harbor seals (*Phoca vitulina*) and gray seals (*Halichoerus grypus*) in Europe in 2016–2021^28–30^ and in North America in May–July 2022^17,19^, the magnitude of those mortalities (<200 deaths in total) would pale in comparison with the impacts that ensued when these viruses arrived in South America. At least 30,000 sea lions have died in Peru, Chile, Argentina, Uruguay and Brazil^10,12–14,20,22–25^. In addition, HPAI caused the largest mortality event of elephant seals recorded to date, with the death of >17,000 pups and an unknown number of adults at Península Valdés, Argentina^26^. Our epidemiological account of this outbreak is the first to provide clinical observations with ecological context for H5N1 HPAI infection in elephant seals. Furthermore, our viral genome data provides evidence for the evolution of a novel marine mammal clade of H5N1 (2.3.4.4b) HPAI virus that has spread among pinnipeds in several countries of South America, revealing mutations that may have enabled their ability to infect mammals while also retaining the ability to spillover to avian hosts.

While serological surveys indicate broad exposure to influenza A viruses (IAV) in pinnipeds globally, mass mortality events have been rare^31–33^. Prior to 2022, the largest IAV outbreak in pinnipeds occurred in 1980, when H7N7 HPAI viruses killed 400–500 harbor seals at Cape Cod, USA, representing ∼20% of the species’ local population^34,35^. Other significant pinniped mortalities attributed to IAV comprise the death of 162 harbor seals in New England, USA, in 2011 due to H3N8 strain^36^ and 152 harbor seals in Denmark in 2014 due to H10N7 strain^37^. Prior to 2023, no pinniped deaths had been attributed to IAV in South America. There are also no published studies reporting on the detection of IAV (or antibodies against them) in southern elephant seals. For northern elephant seals (*Mirounga angustirostris*), the only IAV detections were asymptomatic infections with human-like H1N1 strains in California, USA, in 2009–2012 and 2019^38–40^. Considering that IAV surveys on the Atlantic coast of South America have only reported low pathogenicity avian influenza (LPAI) H11 and H13 strains in coastal birds^41–43^ and antibodies against H1 strains in fur seals^32^, it is likely that southern elephant seals at Península Valdés were naïve to H5 viruses until 2023.

Our data clearly shows that elephant seal pups were severely impacted by H5N1 at Península Valdés, but the extent to which adult elephant seals were affected by HPAI is unclear. The unusually high number of adult carcasses at Punta Delgada, as well as the abnormal haul-outs and the confirmed case in Golfo Nuevo reported here, suggest that adult elephant seals are susceptible to H5N1 (2.3.4.4b) HPAI infection. Furthermore, the complete disruption of the social and breeding structure at Punta Delgada (evidenced by the absence of harems and large alpha males and the presence of motherless pups) suggests that adult elephant seals abandoned the colony prematurely, perhaps after becoming infected. Yet, it is difficult to ascertain the number of adult deaths, which may have happened at sea and will only be accounted for via population censuses at Península Valdés in coming years. Nevertheless, the fact that in 2023 the adult females abandoned the beach before being impregnated (which normally occurs when pups are weaned^44^) suggests that this population will likely experience an atypically low birth rate in 2024, even if most adult females survived.

From a disease evolution standpoint, there is growing concern that H5N1 viruses adapted to mammalian transmission could facilitate host-jumps to other species, including humans. Mammal-to-mammal IAV transmission is believed to have occurred sporadically among pinnipeds over the years^34–36,42,45,46^. The recent demonstration that the H5N1 strain from a human case in Chile (which belongs to the marine mammal clade discussed in this study) is transmissible between co-housed ferrets^47^ also supports the notion that mammal-to-mammal transmission could have played a role in the spread of these viruses in marine mammal communities in South America. We believe that the high mortality rate in elephant seal pups is also consistent with mammal-to-mammal transmission, as pups are toothless and nurtured exclusively through nursing from their mothers. Contact with wild birds is minimal and could not explain the death of ∼95% of all pups born in a matter of weeks. Some newborns may have been infected before birth, as transplacental transmission of H5N1 HPAI viruses has been reported in humans^48^ and high virus loads were detected in aborted sea lion fetuses^10,23^. Yet, how their mothers would have been infected in the first place without mammal-to-mammal transmission presents a thornier question. Feeding is an unlikely route, since the diet of elephant seals relies on squid, fish and crustaceans captured in deep waters^44,49^, and adult elephant seals will fast while on land^44,50^. Moreover, elephant seals are pelagic and only come to shore to breed and later to molt, thus limiting the time window for inter and intraspecific interactions and transmission^44,51^. The main interactions between birds and elephant seals involve opportunistic scavenging of elephant seals’ placental remains, molted skin and carcasses by gulls^50^ (Supplementary Figure 1E), which provides more opportunities for mammal-to-bird transmission than vice-versa. In this context, although there are still many unknowns about the precise viral transmission routes (e.g., contact, environmental, aerosol), mammal-to-mammal transmission seems the most plausible hypothesis to explain the rapid and multinational spread of H5N1 HPAI viruses among pinnipeds in South America.

In the case of Península Valdés, H5N1 infection in sea lions could have been an initial source of virus exposure for the elephant seals. Notably, the epidemic path of HPAI along coastal Argentina left virtually no rookery or stretch of beach without affected sea lions from south to north^23,52^, and then progressed to neighboring Uruguay and Brazil^24,25^. This unrelenting spread along the Atlantic mirrored that seen along the Pacific, with the common denominator being infected sea lions^10,20^ (Figure 5). South American sea lions regularly visit multiple rookeries and haul-outs, sometimes interacting aggressively with other pinnipeds and even killing their pups^53,54^. At Punta Delgada, we observed numerous sea lion carcasses (Figure 1A, Table 2) and witnessed aggressive interactions between sea lions and elephant seals (Supplementary Figures 1C and 1D). Government veterinarians who monitored sea lion rookeries in Argentina noted that animals showing clinical signs of HPAI survived for several days and often abandoned the rookeries while ill (Veronica Sierra, pers. comm.). It is plausible that these sea lions visited different sites during their convalescent period, including elephant seal colonies, and may have played a key role in the spread of H5N1 viruses. In addition, mammal-to-bird spillovers do not seem improbable given the frequent observations of gulls and other avian scavengers feeding on sea lion and elephant seal carcasses in Argentina (Supplementary Figure 1E). It is unclear how terns (which are not scavengers) were infected, and further studies may help clarify the potential role played by gulls as bridge hosts from pinnipeds to other seabirds.

Mammal-to-mammal transmission is also supported by our regional phylogenetic analysis, which identified a novel H5N1 2.3.4.4b clade with viruses that appear to be specific to marine mammals. This marine mammal clade comprises strains with mutations that were not present in H5N1 2.3.4.4b viruses in birds (wild and domestic) from Peru, Chile, Argentina, Uruguay and Brazil, excepting occasional spillovers from marine mammals to coastal birds (terns and sanderling; Supplementary Tables 4 and 5). Some of these mutations (such as Q591K and D701N in PB2) are associated with increased virulence, transmission, or adaptation to mammalian hosts^55–57^ and have been maintained since they first emerged in H5N1 HPAI viruses in marine mammals in Chile. The maintenance of a unique cassette of mutations in viruses from marine mammals (Figure 4), the lower rate of evolution of these viruses (Figure 3B), and the distinct pathways of spread across host groups (Figure 3A) and geographical areas (Figure 5), strongly support the hypothesis that viruses from the novel H5N1 marine mammal clade had an independent chain of virus transmission among marine mammals, separate from the avian transmission chains in Argentina and other countries, and retained the capacity to spillover to terns.

To our knowledge, the H5N1 HPAI 2.3.4.4b marine mammal clade identified in South America represents the first multinational transmission of HPAI in mammals ever observed globally. Over the last century, LPAI H1, H2, H3, and H7 viruses have periodically jump into mammals, including humans, swine, canines, and equines, causing major outbreaks and pandemics^58^. Spillover of H5N1 2.3.4.4b clade also regularly occurs in humans and terrestrial and marine mammals on a global scale, but onward transmission in mammals is limited and not sustained over time, leading to speculation that the H5 subtype perhaps is not capable of causing a pandemic^19,59,60^. Despite gaps in the available data, our epidemiological and phylogenetic results support the hypothesis that the spread of viruses from the novel marine mammal clade in South America has occurred via mammal-to-mammal transmission, but as with any hypothesis, this is subject to revision as more data become available.

The implications of sustained mammal-to-mammal transmission of H5N1 HPAI viruses could be far-reaching, both from a conservation and a public health perspective. From the standpoint of wildlife conservation, this is particularly concerning for endangered pinnipeds with limited geographic distribution such as Caspian seals (*Pusa caspica*), Hawaiian monk seals (*Neomonachus schauinslandi*), among others^61^. Significant mortalities of southern elephant seals and Antarctic fur seals have already been attributed to H5N1 HPAI in South Georgia^62,63^, although it is not clear whether the viruses involved in these cases belong to the marine mammal clade identified in this study or if instead they are similar to viruses detected in brown skuas (*Stercorarius antarcticus*) from the same archipelago, which cluster with avian viruses from inland Argentina^62^. The detection of marine mammal clade viruses in dead dolphins and porpoises in Chile^14^ is also concerning, since 23% of the world’s odontocete species are already threatened with extinction^64^. If pinnipeds become a sustainable reservoir for H5N1 HPAI viruses that retain the capacity to infect wild birds, coastal bird species could be repeatedly affected by spillover infections. Furthermore, the implications could become even more severe if the marine mammal clade viruses evolve to enable transmission among terrestrial mammals, or if additional gene reassortment occurs with South American LPAI viruses present in Argentina^41,43,65,66^, potentially expanding either the host range, pathogenesis and/or transmission in wildlife.

From a public health perspective, mammal-to-mammal transmission could be a critical stepping-stone in the evolutionary pathway for these viruses to become capable of human-to-human transmission and thus potentially pandemic^67^. As mentioned previously, some of the mutations found in the strains of the marine mammal clade are already known to be of concern. Specifically, the mutation D701N in PB2 has been shown to compensate for the lack of the E627K mutation in PB2 in terms of improved viral growth in mammalian cells and enhanced aerosol transmission of H3N2 and H5N1 viruses^68^. On the other hand, the phenotypic effects of mutations in other gene segments found in the H5 viruses from our study (Supplementary Table 5) are not yet known, and the possibility that some of them may also open evolutionary pathways that enhance the virulence or transmission of these viruses to mammals (including humans) cannot be ruled out. The fact that the H5N1 HPAI virus detected in a human case in Chile^69^ belongs to the marine mammal clade described in this study, highlights the potential risk to public health. Moreover, given pinniped susceptibility to multiple IAVs (including human-like strains^38–40^), and their frequent intermingling with other avian and mammalian hosts, co-infections could occur, potentially enabling the emergence of reassorted strains^36,70^. Hence, while there is no evidence for genomic reassortment occurring in pinnipeds at this time, the broad circulation of H5N1 HPAI viruses in marine mammals is a warning we must not ignore.

In conclusion, as recently demonstrated by the detection of HPAI H5N1 viruses in ruminants^71^, few if any compartments and species are outside the scope of the clade 2.3.4.4b strains. Thus, moving forward, HPAI management requires holistic strategies that recognize the interconnectedness of human, animal, and environmental health and safeguard biodiversity, promote sustainable practices, and enhance resilience globally to emerging infectious diseases.

## METHODS

### Study species

Southern elephant seals are widely distributed in Subantarctic islands, with a single continental colony at Península Valdés, Patagonia, Argentina (representing ∼5% of the global population)^61^. The species has a well-defined annual life cycle, which we summarize as follows based on studies at Península Valdés^44,51^. Adult (and subadult) males and females haul-out in late August and early September, with alpha males establishing and defending harems (median 11–13 females per harem, with a maximum of 134 females); subordinate males are chased away but remain along the margins of harems.

Most females are pregnant when they come ashore, giving birth within 5.7±1.9 days after arrival (80% of pups are born by 2 October). Pups are toothless and will nurse for 22.4±1.7 days; during this period the females will fast and remain with their pups, under the protection of the alpha male. Copulations will begin 20.3±2.1 days after parturition, i.e. shortly before females wean their pups. The female then abandons the pup and returns to the sea to forage; on average, females spend a total of 28.2±2.5 days ashore, fasting. Males also fast on land and will abandon the beach approximately at the same time as females; adult seals are nearly absent by mid-November. Weaned pups will remain on the beach for several weeks, fasting while they complete their development and are ready to go to sea to forage. Juveniles and adults will return to the beaches later in the season to undergo molt, with juveniles molting earlier (November to January) than subadults and adults (December to February).

### Study site and field observations

Península Valdés is located in Chubut, Argentina, and is a UNESCO World Heritage site of global significance for the conservation of marine wildlife. We studied two sites at Península Valdés: the elephant seal breeding colony at Punta Delgada and the interior beaches of Golfo Nuevo where sporadic seal haul-outs occur. Punta Delgada (from 42.753°S 63.632°W to 42.771°S 63.649°W) is a 3-km beach on the exposed seashore of Península Valdés (Supplementary Figure 1) where southern elephant seals breed in high densities^72,73^. Field surveys were conducted on 5-Oct-2022 (baseline year), and during the mortality event on 10-Oct-2023, 3-Nov-2023 and 13-Nov-2023. In each survey, a team equipped with binoculars walked along the clifftop to count live and dead elephant seals, differentiating individuals by sex and age class (pup, weaner, juvenile, subadult male class 1–4, adult male, adult female) and male dominance status (alpha or subordinate)^74,75^. For outbreak investigation in 2023, a second team of trained veterinarians wearing full PPE descended to the beach to document clinical signs and collect samples from affected animals and count the carcasses of other wildlife species. We also covered a 50-km stretch of interior beach in Golfo Nuevo, from Cerro Prismático (42.595°S 64.811°W) to Cerro Avanzado (42.835°S 64.874°W), including the city of Puerto Madryn (∼130,000 inhabitants) (Supplementary Figure 2). Elephant seals do not breed in this area, but sporadic haul-outs are reported by the public and park rangers to the Red de Fauna Costera de la Provincia del Chubut (RFC). Data on the age, sex, condition, location, and date of each seal were extracted from RFC records for 2022 and 2023.

### Sample collection

On 10-Oct-2023, a team of trained veterinarians wearing full PPE descended to the beach at Punta Delgada to collect samples from affected animals. Post-mortem swabs (oronasal, rectal, tracheal, lung and brain) were collected from four elephant seal pups, six South American terns (*Sterna hirundinacea*) and two royal terns (*Thalasseus maximus*) found dead (carcasses still in *rigor mortis*). On 1-Nov-2023, swabs were obtained from a subadult male elephant seal that hauled-out and died in Golfo Nuevo. Swabs were placed in cryotubes containing 1 mL of DNA/RNA Shield (Zymo Research, Irvine, CA, USA) for inactivation, and stored in a cooler with icepacks, then transferred to –80°C within 24 hours.

### Virus detection

Samples from four elephant seal pups, five adult South American terns and two adult royal terns were pooled according to species and sampled tissue and other pools were prepared with all samples from the dead subadult male elephant seal (oronasal, rectal and lung) and from a juvenile South American tern (brain, lung, oronasal and rectal). Viral RNA was extracted from 140 µL of suspension from swabs using a QIAamp Viral RNA Mini Kit (Qiagen, Valencia, CA, USA). RNA was eluted in a final volume of 60 µL and stored at –80°C. Viral cDNA was prepared using 15 µL of viral RNA and random hexamers in a final volume of 30 µL using a High-Capacity cDNA Archive kit (Applied Biosystems, Foster City, CA, USA). The cDNA from all pooled samples were tested for influenza A viruses by RT-qPCR using TaqMan Universal PCR Master Mix (Applied Biosystems) directed to the matrix gene^76^. Positive samples from elephant seals were then tested using primers and probes for H5 clade 2.3.4.4b detection^77^. Quantification cycle (Cq) values were used as a proxy to compare viral RNA load in different samples and to facilitate sample selection for full genome sequencing. RT-qPCR reactions were performed on an ABI Prism 7500 SDS (Applied Biosystems).

### Full genome sequencing

The viral genome was amplified from RNA using a multi-segment one-step RT-PCR with Superscript III high-fidelity RT-PCR kit (Invitrogen, Carlsbad CA) according to manufacturer’s instructions using the Opti1 primer set (Opti1-F1, Opti1-F2 and Opti1-R1) previously described^78^. Amplicons were visualized on a 1% agarose gel and purified with Agencourt AMPure XP beads (Beckman Coulter, Brea, CA). The concentration of purified amplicons was quantified using the Qubit High Sensitivity dsDNA kit and a Qubit Fluorometer (Invitrogen). The sequencing library was prepared with the Rapid Barcode library kit SQH-RBK110.96 (Oxford Nanopore, Oxford, UK) and loaded on the Mk1c sequencer according to ONT instructions for the R.9 flow cells. Real time basecalling was performed with MinIT (Oxford Nanopore); the automatic real time division into passed and failed reads were used as a quality check, excluding reads with quality score < 7. Quality-checked reads were demultiplexed and trimmed for adapters and primers, followed by mappings and a final consensus production with CLC Genomics Workbench v23.0.2 (Qiagen).

### Phylogenetic analysis

To place the coastal Argentinean viruses in a global context, we downloaded HA gene sequences from HPAI H5N1 clade 2.3.4.4b viruses globally from GenBank and GISAID since January 1, 2021. Phylogenetic relationships were inferred for HA gene using the Maximum likelihood (ML) methods available in IQ-Tree 2^79^ with a GTR model of nucleotide substitution with gamma distributed rate variation among sites. Due to the size of the dataset, we used the high-performance computational capabilities of the Biowulf Linux cluster at the National Institutes of Health (http://biowulf.nih.gov). To assess the robustness of each node, a bootstrap resampling process was performed with 1000 replicates.

To study how the H5N1 HPAI outbreaks in Argentina were connected to outbreaks occurring in other South American countries, we performed a phylogenetic analysis of 11 available H5N1 virus genomes from Patagonia Argentina from three species of marine mammals and two species of terns, along with 225 closely related H5N1 virus genomes obtained from avian and mammalian hosts in five South American countries (Argentina, Brazil, Chile, Peru, Uruguay) and Antarctica available from GISAID and/or GenBank public databases (Supplementary File 2). Alignments were generated for each of the eight segments of the virus genome (PB2, PB1, PA, HA, NP, NA, MP, and NS) using MAFFT v7.490^80^. Phylogenetic trees were inferred for each segment individually using maximum-likelihood methods with a GTR+G model of nucleotide substitution and 500 bootstrap replicates, using the CLC Genomics Workbench v23.0.2 (Qiagen) and the inferred trees were visualized. Since the H5N1 viruses were collected from a recent outbreak and had little time to accrue mutations and diversify, limiting genetic diversity, all Bayesian analyses were performed using concatenated genome sequences (13,140 nt) to improve phylogenetic resolution (after removing reassortants and viruses that did not have sequence data available for all eight segments).

We performed a time-scaled Bayesian analysis using the Markov chain Monte Carlo (MCMC) method available using the BEAST package pre-release v1.10.5 (compiled on 24-Apr-2023)^81^, using GPUs available from the NIH Biowulf Linux cluster. First, the analysis was run with an exponential growth demographic model, a GTR+G model of nucleotide substitution, and an uncorrelated lognormal relaxed molecular clock. To account for the possibility that high rates of convergent evolution involving adaptive mutations following host-switches (see mutation analysis below) could artificially cluster marine mammal viruses on the tree that do not actually share common ancestry, a second tree was inferred for the third codon position only. The MCMC chain was run separately 3–5 times for each dataset using the BEAGLE 3 library^82^ to improve computational performance, until all parameters reached convergence, as assessed visually using Tracer version 1.7.2^83^. At least 10% of the chain was removed as burn-in, and runs for the same dataset were combined using LogCombiner v1.10.4. An MCC tree was summarized using TreeAnnotator v1.10.4. All XMLs and output files are available in Supplementary File 2.

After the initial analysis determined that the vast majority of H5N1 viruses collected from marine mammals clustered together in a well-supported clade (posterior probability = 1.0), in both the whole genome and third codon analyses, we repeated the BEAST analysis using a more appropriate host-specific local clock (HSLC)^84^ to accommodate differences in the evolutionary rate between marine mammals and avian hosts. For the HSLC analysis, any singleton avian and human viruses positioned in the marine mammal clade (likely representing transient dead-end spillovers) were excluded to ensure monophyly. Similarly, any singleton marine mammal viruses positioned in the major avian clade (which also likely represent transient dead-end spillovers from birds to marine mammals) were excluded.

To compare evolutionary rates in marine mammals and avian hosts across the eight different segments of the virus genome, the analysis was repeated using eight genome partitions (PB2, PB1, PA, HA, NP, NA, MP, NS). A phylogeographic discrete trait analysis^85^ was performed to quantify rates of viral gene flow between different host groups (wild bird, poultry, marine mammal, human) as well as between locations (Argentina, Brazil, Peru, Chile, Uruguay, Antarctica). Since extensive virus gene flow was observed between Chile/Peru, which is not the focus of this study, a single combined Chile/Peru location category was used. A location state was specified for each viral sequence based on the host species and location of collection. The expected number of location state transitions in the ancestral history conditional on the data observed at the tree tips was estimated using ‘Markov jump’ counts^86,87^, which provide a quantitative measure of asymmetry in gene flow between defined populations. To estimate absolute rates of synonymous and non-synonymous substitutions as well as dN/dS, we employ a ‘renaissance counting’ procedure that combines Markov jump counting with empirical Bayes modeling^88^. R v4.3.2^89^ was used to summarize and visualize the outputs of these analyses.

### Mutation analysis

Consensus nucleotide sequences for the eight open reading frames were translated to protein and compared to viruses from birds and mammals from Argentina, other South American countries, Antarctica, North America (genotype B3.2 from 2022–2023), and reference strains from Asia (A/goose/Guangdong/1/1996 and A/Vietnam/1203/2004).

## DATA AVAILABILITY

We gratefully acknowledge the authors and both originating and submitting laboratories of the sequences from GISAID’s EpiFlu™ Database on which this research is based. GenBank accession numbers for all the sequences generated as part of this study are provided in Supplementary Table 3. In addition, XMLs, MCC and ML trees, and GISAID acknowledgement tables are also provided in Supplementary File 2.

## ACKNOWLEDGEMENTS

Wildlife Conservation Society, University of California, Davis and the National Institute of Agricultural Technology (INTA) (PNSA PD114) funded this study. This work was also partially supported by Alexander von Humboldt Foundation, through a fellowship to A. Rimondi. We thank Lic. M. Cabrera from Dirección de Conservación, Secretaría de Turismo Municipal de Puerto Madryn and park rangers from Área Natural Protegida El Doradillo. We also thank H. O. Loza from Reserva Natural de la Defensa Faro Punta Delgada (Armada de la República Argentina) and the navy personnel that work there. We acknowledge data shared by Red de Fauna Costera de la Provincia de Chubut. Permits were granted by Dirección de Fauna y Flora Silvestres de la Provincia de Chubut and Subsecretaría de Conservación y Áreas Protegidas de Chubut. We acknowledge Servicio Nacional de Sanidad y Calidad Agroalimentaria (SENASA) and other laboratories in South America that submitted H5N1 virus sequence data to GISAID.

## AUTHOR CONTRIBUTIONS

Author contributions were as follows: Study design: M.M.U., A.R. Funding: M.M.U., V.F., A.R. Sample and data collection: M.M.U., R.E.T.V., J.C., V.Z., V.F., C.C. Virus detection and virus sequencing: V.S.O., A.R. Phylogenetic analyses: M.I.N., A.R., P.L. Data analysis and interpretation: M.M.U., R.E.T.V, M.I.N., A.R. Writing of the manuscript: M.M.U., R.E.T.V., M.I.N., A.R. All authors approved the manuscript before submission.

